# Non-uniform chromosomal SNP density biases sites of meiotic crossovers in *Drosophila melanogaster*

**DOI:** 10.64898/2025.12.30.697082

**Authors:** Savanna Hinson, Ryan Sangston, Karol Cichewicz, Guruprasad Konduru, Ishan Parikh, Jay Hirsh

## Abstract

Here we localize a genetic suppressor that enhances the reduced locomotor activity phenotype of flies lacking brain dopamine. We utilized a non-bulked segregant analysis using whole genome sequencing (WGS), mapping the trait to a roughly 3.5 mega-base (Mb) region of the X-chromosome. However, this mapping yielded ∼5-fold lower resolution than anticipated, due to an uneven distribution of Single Nucleotide Polymorphisms (SNPs) between the X-chromosomes of the two recombining lines. This uneven SNP distribution was associated with recombination events biased towards regions of low SNP density, and away from the more SNP dense regions associating with the activity phenotype. We find that nearly perfect mapping of X-chromosome visible markers occurs only in historical data from a time before the establishment of discrete genetic background strains. This suggests that genetic uniformity in early *Drosophila* studies may have contributed to more consistent recombination frequencies, whereas modern mapping efforts are complicated by variability in SNP distribution across recombining strains. These findings highlight challenges in *Drosophila* genetic mapping in situations where altered SNP density can skew recombination, complicating trait localization.

**Article Summary:** Genetic mapping studies generally assume uniform crossover distribution across chromosomes. However, this study demonstrates that an uneven density of single nucleotide polymorphism (SNP) is associated with biased sites of meiotic recombination. Using brain dopamine-deficient *Drosophila*, SNP dense regions show reduced crossover frequency, which reduced the mapping resolution, complicating genetic trait localization. These findings highlight the need to consider parental chromosome SNP distribution and its impact on recombination when designing genetic mapping studies. Future studies should consider chromosome structure, parental haplotype, and sequence heterogeneity to enhance mapping accuracy and resolution.

## Introduction

Genetic mapping enables the identification of loci associated with phenotypic traits, and the expected resolution of genetic mapping relies on the simplified assumption that meiotic crossovers (COs) are randomly distributed across the genome. However, deviations from this assumption have been observed in many species, including *Drosophila melanogaster*. Studies in *Drosophila* have shown variation in recombination rates across chromosomes, with regions near centromeres and telomeres suppressing crossovers and significant variation also within the interior of a chromosome (Lindsley & Sandler, 1977), (Chan et al., 2012; Comeron et al., 2012; Hughes et al., 2018).

Here, we examine how variations in parental SNP density alter crossover frequency and locations in recombinant offspring. Our work began with an investigation into the genetic basis of locomotor recovery in dopamine-deficient flies. Dopamine (DA) has two critical functions in both the central nervous system (CNS) and cuticular processes in *Drosophila*. The biosynthesis of dopamine is regulated by tyrosine hydroxylase (TH), encoded by the *pale* (*ple*) gene, which generates distinct splice variants for CNS and cuticular functions (Birman et al., 1994). By restoring cuticular dopamine synthesis in a *ple*-lethal background, we generated viable flies that are entirely devoid of CNS dopamine (Cichewicz et al., 2017; Riemensperger et al., 2011). These flies initially exhibited reduced locomotor activity, consistent with previous findings showing that dopamine is required for normal behaviors (Friggi-Grelin et al., 2003; Lima & Miesenböck, 2005; Yellman et al., 1997). While ‘cleaning up’ the genetic background in this line, we discover a genetic suppressor of the hypoactivity in these flies, leading to near normal levels of locomotion, with no restoration of DA.

To identify the genetic factors responsible for this unexpected recovery of locomotor function, we employ a non-bulked segregant analysis using whole genome sequencing (WGS). Our analysis maps the trait to a 3.5 megabase (Mb) region on the X chromosome, a substantially lower resolution than expected. We identify an uneven distribution of SNPs within the X chromosomes of the two parental lines, which is associated with a bias in crossover locations and frequency. Regions of near homozygosity, or low SNP density, exhibit higher crossover frequencies, while regions with higher SNP density show reduced crossovers.

## Results

A viable line of flies wholly lacking in brain dopamine was isolated by Cichewicz et al, 2017, by rescuing a lethal mutation in the *pale (ple)* gene with a 3rd chromosome bacterial artificial chromosome (BAC) that selectively rescues *ple^+^* function in the hypoderm, but not in CNS. As expected, these flies show reduced levels of locomotor activity relative to wild type controls. In the process of ‘cleaning up’ the genetic background of these flies, they were crossed to our laboratory stock of *w^1118^*, and the relevant 3rd chromosomes were reisolated by single pair matings. To our surprise, a small number of the stocks resulting from the single pair matings show normal levels of locomotor activity. Activity levels of one of these stocks, which we name *DD-Hi*, is shown in Figure 1, compared to our lab *w^1118^* and to stocks showing the more expected and reduced levels of activity, hereafter named *DD-Lo*.

**Figure 1.**
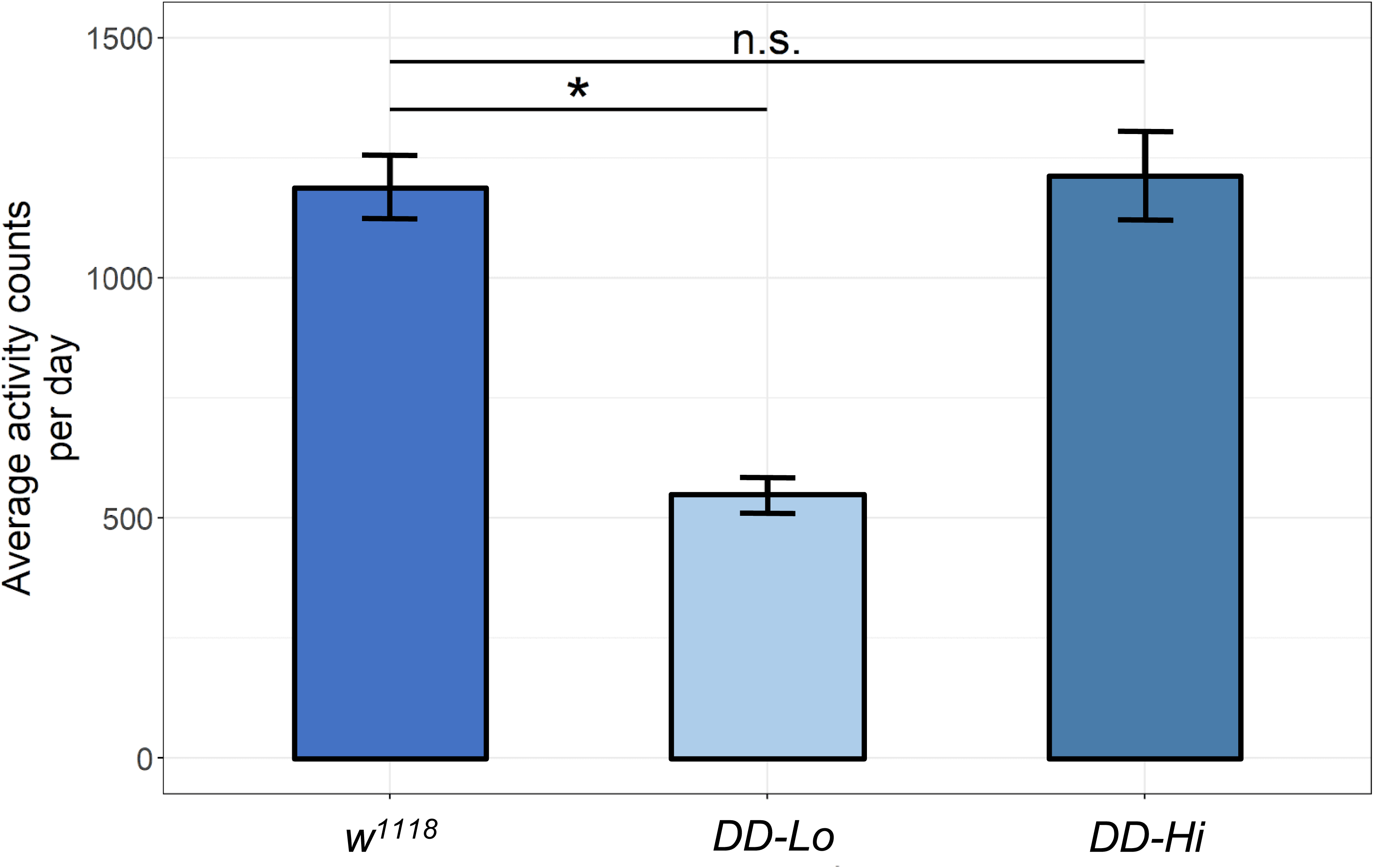
Activity levels of the flies used in this study. Average activity levels in constant darkness in our lab stock of *w^1118^*, *DD-Lo* and *DD-Hi*. 3–5-day old adult flies were entrained to 12:12 LD cycles for 5 days, then switched to constant darkness (DD). Activity levels were averaged over 3 days in DD. Significance indicates p<0.05 tested by an ANOVA followed by Tukey’s honest test of significance.

Given the surprisingly high levels of locomotor activity levels in the *DD-Hi* line, we sought to validate that these flies were still deficient in brain dopamine. We did this using two approaches, first measuring DA in brain extracts by HPLC with electrochemical detection (Hardie & Hirsh, 2006), and second, by probing these flies pharmacologically. Both of the *DD* lines show no detectable DA by HPLC analysis, whereas serotonin levels are unaffected (Figure 2A). Since this approach could miss trace levels of brain DA, we followed up with a pharmacologic approach. Cichewicz et al (2017) found that locomotor activity of the initial stock of brain dopamine deficient flies was insensitive to feeding of the tyrosine hydroxylase inhibitor, 3-IY. The *DD-Lo* and *DD-Hi* lines are similarly insensitive to 3-IY, whereas the *w^1118^* control line shows reduced locomotor activity after 3-IY feeding (Figure 2B). In addition, we fed the flies methamphetamine, which stimulates dopamine release and blocks reuptake (Nestler, 2004), thus stimulating locomotor activity in adult *Drosophila* (Andretic et al., 2005). The control *w^1118^* line shows greatly enhanced locomotion after methamphetamine feeding, but the locomotor activity of the *DD-Hi* or *DD-Lo* lines are unaffected (Figure 2B). These results indicate that the *DD-Hi* flies have regained normal levels of locomotor activity in a manner that is independent of even trace levels of neural dopamine.

**Figure 2.**
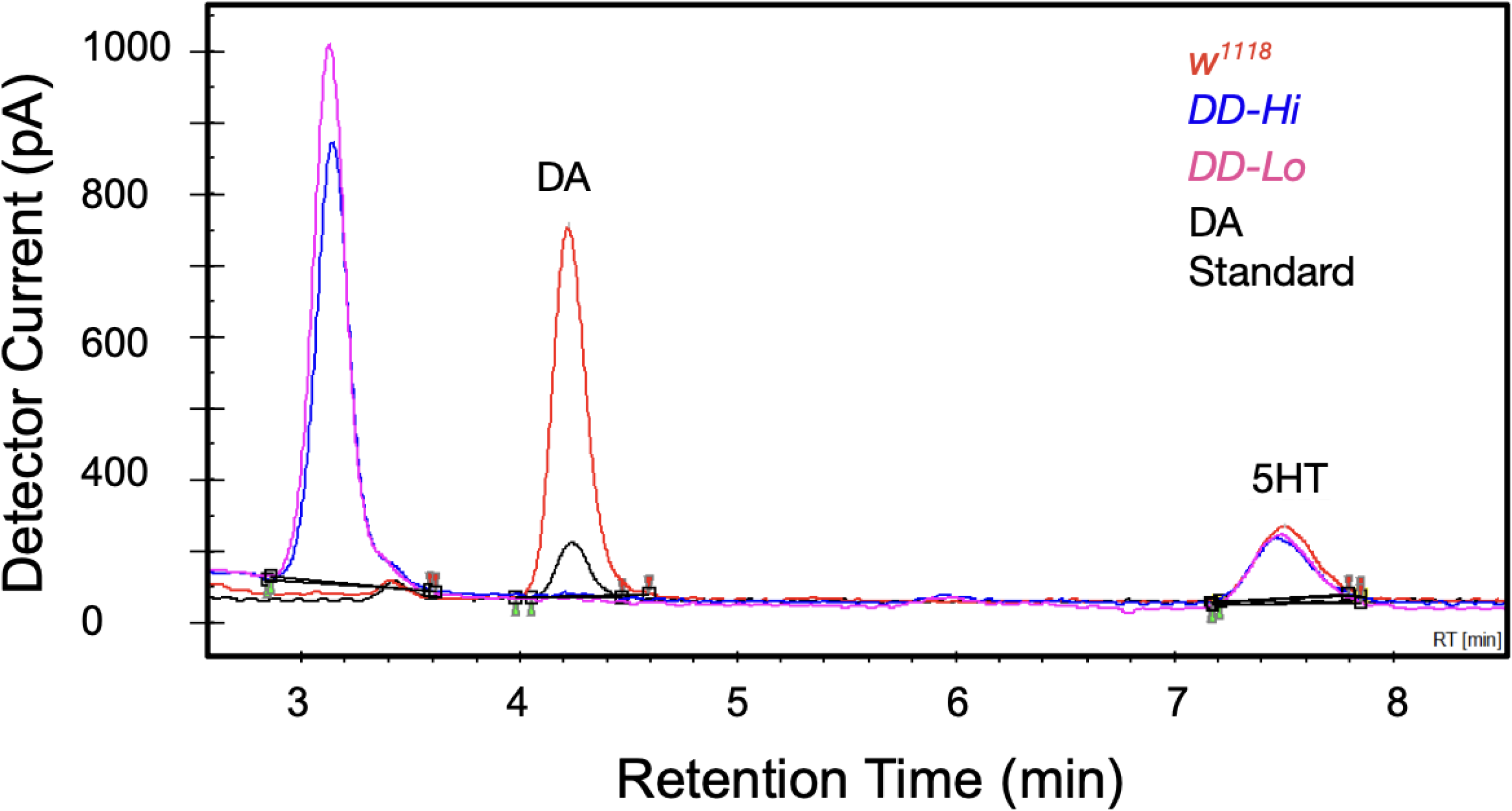
Dopamine deficient lines lack DA by HPLC or via pharmacologic intervention. A) Neither *DD-Lo* or *DD-Hi* brain extracts show detectable DA. A peak with the retention time characteristic of DA was detected in the standard and in brain extracts from *w^1118,^* but no DA peak was detected in brains of *DD-Lo* or *DD-Hi.* Normal levels of 5-HT were detected in all three genotypes. The peak with the retention time of 3.1 min is associated with red eye pigment and was detected in the two DA-deficient genotypes, which express the *white^+^* transgene. Y axis scale: 1V= 2nA detector current. B) Locomotor activity analysis of flies treated with the TH inhibitor 3-iodotyrosine (3-IY), and methamphetamine (METH), demonstrates a DA-independent mechanism responsible for hyperactivity of *DD-Hi* line. Locomotor activity was normalized to No Drug control for each genotype. [*p<5E−04, one-way ANOVA (Bonferroni-corrected), error bars represent SEM, n=22 to 32].

### Identification of the gene(s) responsible for the altered locomotor activity in *DD-Lo* vs *DD-Hi*

We sought to identify the genetic element(s) responsible for the unexpected *DD-Hi* phenotype via non-bulked segregant analysis with WGS (Michelmore et al., 1991). In this study, the *DD-Hi* and *DD-Lo* stocks were allowed to freely intermate for fifteen generations, followed by measurement of locomotor activity in individual recombinant flies, then identifying the single nucleotide polymorphisms, SNPs, that associate significantly with activity levels. The starting *DD-Hi* and *DD-Lo* stocks showed a high level of isogeny on the X chromosome, with 99.93% homozygosity for *DD-Hi* and 99.95% for *DD-Lo*. Regions of the *DD-Hi* and *DD-Lo* parental chromosomes that contained variation among samples were removed from our analysis (7.9% and 5.8%) of SNPs were heterozygous, respectively). We thus focused solely on homozygous SNPs within each parental line to eliminate any confusion over non-fixed SNPs.

Association was performed by measuring activity levels in ∼800 recombinant male flies (see Methods), then dividing these into two pools representing the high and low extremes of activity levels. The 15-generation recombinant male flies were separated into two bulked groups of 45 flies each, a low activity group with activity levels from 95 to 238 counts/day, mean=194, and a high activity group with 1460 to 1861 counts/day, mean=1609. Figure 3A shows the results of this mapping study, showing SNPs with significant association on the X chromosome mapping to a ∼3.5 Mb region of the X-chromosome from 14 to 18 Mb, with a gap at ∼16.5-17 Mb, lower resolution than we had hoped for. Since recombination occurs only in female *Drosophila*, and females contain two X-chromosomes vs the single X-chromosome in males, a given X-chromosome will undergo on average 10 generations of meiosis in females, and given that only half of the chromatids will have one crossover in each meiosis, there will be 5 opportunities for single reciprocal recombination events (Jones et al., 2024).

**Figure 3.**
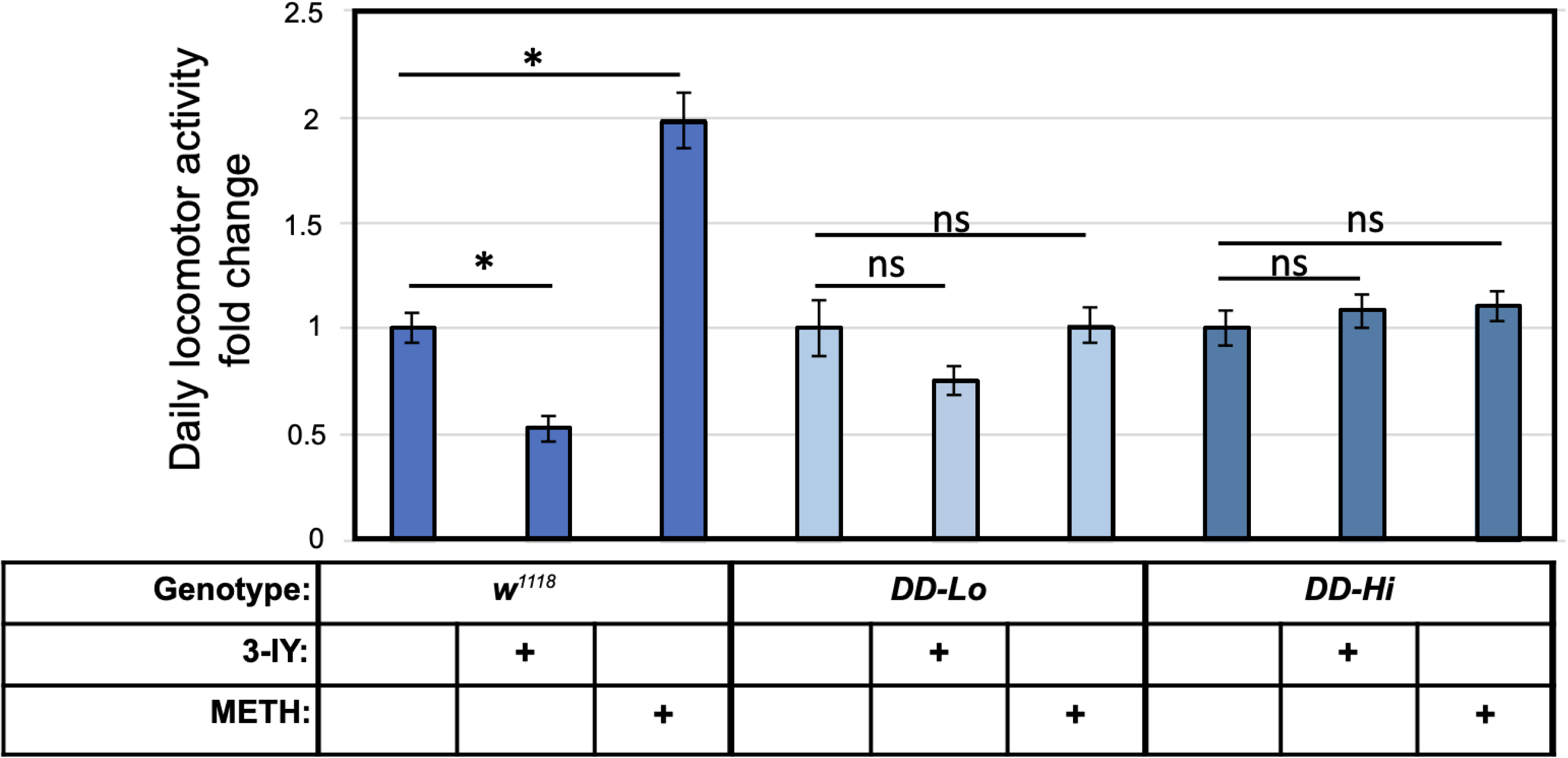
A non-bulked segregant analysis localizes the SNPs associating with high vs low locomotor activity after 15 generations free recombination between the *DD-Hi* and *DD-Lo* lines, and *DD-Hi* and *DD-Lo* X-chromosomes show unevenly distributed single nucleotide differences. A) SNPs were determined using the GATK pipeline (Methods). Significantly associating SNPs were determined as p<0.05 after Bonferroni correction, as shown by the black horizontal line. B) Non-uniform distribution of X-chromosome SNPs between the *DD-Lo* and *DD-Hi* X-chromosomes. SNP density is plotted as SNPs/kilobase, with SNPs determined by GATK. The average number of SNPs within the SNP dense regions/10kb are 23 for *DD-Lo* and 27 for *DD-Hi.* The regions that show association with the DD Hi/Low bulked phenotypic differences from the 15-generation non-bulked segregant analysis are shown highlighted in yellow. These regions are exclusively within the regions of enhanced SNP density. The SNP dense regions are labeled A through E in black at the top of the figure. [n=40 high activity and 44 low activity recombinants]

This result was disappointing, since if there is on average one crossover per meiosis per X-chromosome (Miller et al., 2012) and these were randomly distributed, we should have achieved far better resolution, with one crossover on average every 0.74 Mb (23.5 Mb/2^5^). We chose to ignore the autosomes for several reasons; the high activity phenotype occurs only in males and mapped predominantly to the X chromosome, as well as the difficulty in analyzing diploid 2^nd^ and 3^rd^ chromosomes.

There are several possible explanations for this failure to tightly resolve the SNPs associating with the locomotor activity differences between *DD-Hi* and *DD-Lo*. First, an X-chromosome inversion could be suppressing local recombination frequency. Second, there could be a bias in where recombination is occurring, from an uneven distribution of SNPs across the X-chromosome. Third, the entire associating region may be required for the altered locomotor activity between *DD-Hi* and *DD-Lo* phenotype, with many loci acting additively. We investigated each of these possibilities.

### Could inversions be affecting X-chromosome recombination frequency?

The presence of chromosomal inversions could be an additional factor suppressing homologous recombination (reviewed in (Hoffmann & Rieseberg, 2008)), but we didn’t find any after utilizing long read sequencing. We sequenced *DD-Hi* and *DD-Lo* using long read Nanopore sequencing, then used the tool SVIM for analysis (see Methods). No inversions were found that were strongly supported by the data.

### SNP Frequency and Distribution

To investigate the possibility that the SNP frequency/distribution could be biasing sites of recombination, we determined single nucleotide differences in *DD-Hi* vs *DD-Lo,* as shown in Figure 3B. This figure shows a very uneven pattern of SNPs in both *DD-Hi* and *DD-Lo* parental X chromosomes that are clustered into five discrete regions averaging 25 SNPs/10 kb, or 0.25% (Figure 3B) versus 4 SNPs/10kb in the SNP sparse regions, or 0.04%. It is of interest that two of these SNP dense regions overlap with the regions shown in Figure 3A to associate with the differences in locomotor activity between *DD-Hi* and *DD-Lo,* regions that are highlighted in yellow in Figure 3B. Since homologous pairing is a general prerequisite for recombination (Mao et al., 2008), this uneven distribution of SNPs may bias recombination break points to the highly conserved, or SNP sparse regions.

To test the possibility that reciprocal crossovers are affected by this uneven distribution of SNPs, we mapped recombination points on the X-chromosome from the parental females in Figure 3B and classified them relative to the SNP-dense regions, as shown in Figure 4. SNPs called by the GATK short variant discovery pipeline (https://gatk.broadinstitute.org/hc/en-us) were analyzed for parental origin, either from *DD-Hi* (dark blue) or *DD-Lo* parents (light blue), or ambiguous regions (white), which contain non-informative regions of the chromosome or missing sequencing reads. Each row corresponds to an individual recombinant fly, ordered by phenotype from highest activity at the top and lowest activity at the bottom. The middle black line indicates the separation between recombinant samples used for the high activity versus low activity bulked phenotype for association testing, where low activity flies had daily activity counts of 95-238, and high, 1460-1861. The five SNP dense regions are shown in black at the top of the figure, labeled A-E. Figure 4 indicates the importance of SNP dense regions C and D to the phenotypic differences in the recombinant flies, and that reduced CO frequency and randomness of COs both contribute to the lower than expected resolution of our genetic mapping study above.

**Figure 4.**
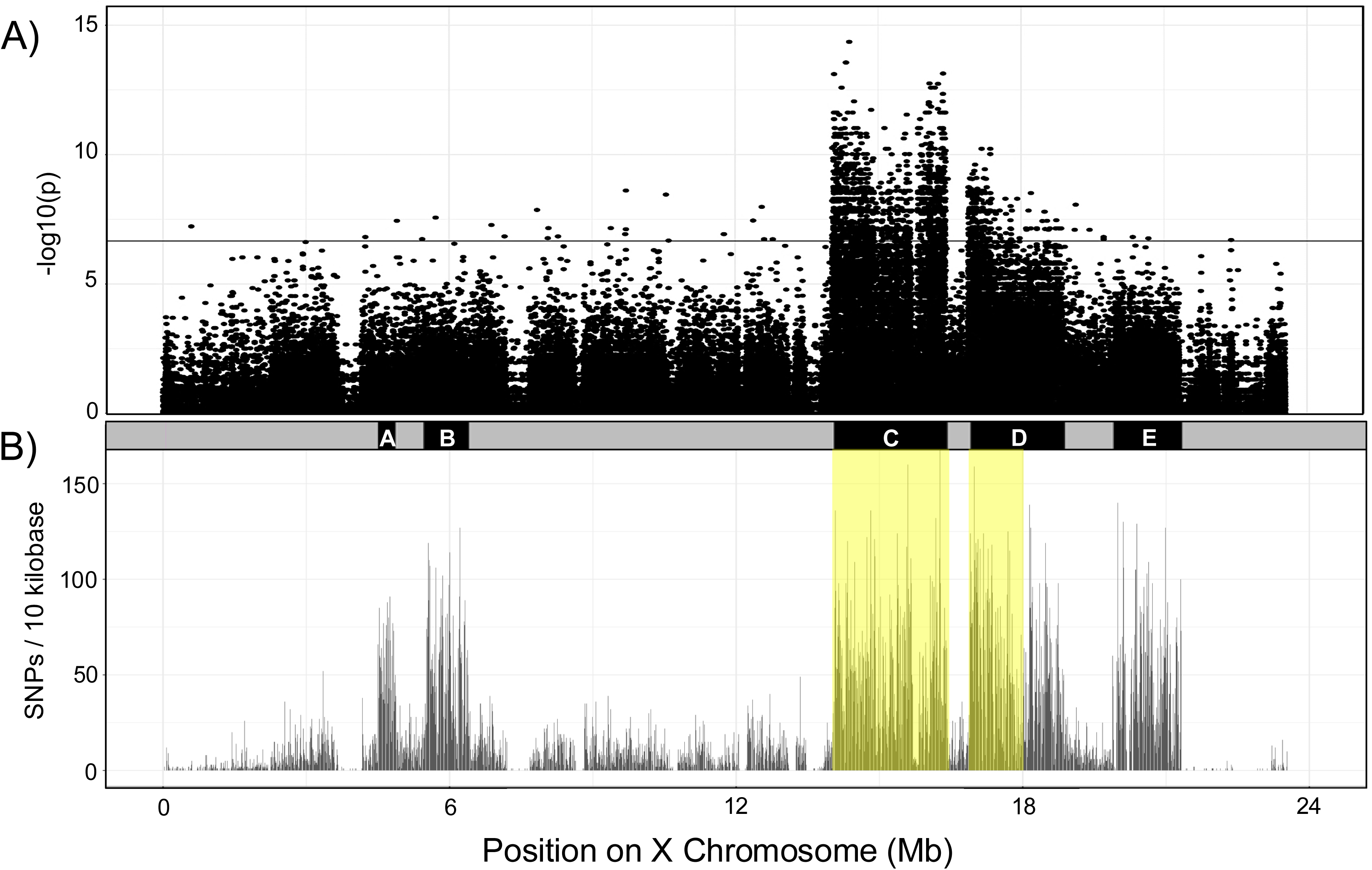
Single nucleotide differences of each individual recombinant fly from the non-bulked segregant analysis and an expanded view of the 0.1 Mb crossover hotspot at 14.15-14.25 Mb. A) Each horizontal line shows one sequenced individual fly, with activity levels ordered by activity from highest activity at the top to lowest at the bottom, with the horizontal line in the center dividing the high vs low bulked flies. SNPs for each sequenced individual were called using the GATK pipeline. Informative SNPs indicating parental lineage were identified by searching for locations where all SNPs called were (1) the same across three parental sequenced samples and differed and (2) different between *DD-Hi* and *DD-Lo*. Informative SNPs were then used to determine if a region matched *DD-Hi* (dark blue), *DD-Lo* (light blue), reference (white), unknown, or possessed a unique (gray) SNP. The SNP dense regions A-E are shown in black at the top of the figure, with the yellow bar in regions C and D indicating the regions showing significant association in the GWAS. Reciprocal recombination events within the SNP dense regions are determined by a switch in parentage from *DD-Hi* to *DD-Lo,* or vice versa, and are indicated by a yellow triangle. The smallest SNP sparse region is indicated with a purple box at the top of the figure.

### Distinct haplotype blocks identified in SNP dense regions

Large regions of unbroken parentage, or haplotype blocks, are apparent in the SNP dense regions, but not in the SNP sparse regions (Figure 4A). The latter is at least partly due to the difficulty of determining parentage in the highly similar SNP sparse regions. The finding of these long haplotype blocks is consistent with failure to better resolve the associating regions. Note that these haplotype blocks are on a faint background of the apparently opposite parentage. We cannot distinguish whether this background represents non-crossover conversion events, versus sequence errors/ambiguity resulting from the marginal sequence coverage from this single recombinant fly sequencing.

### Reciprocal crossovers are reduced in SNP dense relative to SNP sparse regions

A second factor in the reduced resolution of our sequencing is that the frequency of reciprocal crossovers is disfavored in the SNP dense regions relative to an adjacent SNP sparse region. As shown above, COs were localized in the SNP dense regions by a change in haplotype parentage tracts, indicated by yellow triangles in Figure 4. We compared the observed CO frequencies of SNP dense regions A-E with the smallest SNP sparse region between SNP dense regions C and D (shown in purple in Figures 4A and 5). Directly counting COs in the SNP sparse region isn’t possible since we can’t discern haplotype parentage blocks due to the near-identity of parental sequences. Instead, we infer that at least one CO has occurred due to change in parentage in the SNP dense regions on either side of the SNP sparse region, as shown by purple bars in this region in each recombinant fly. The observed rate of inferred COs will be an absolute minimum, since we can’t detect COs if an even number of COs occurs, and our estimated frequency will be reduced if there is selection for the same parentage in the adjoining regions SNP dense regions C & D, as we demonstrate below. Even with this minimum estimate, we observe a recombination frequency that is 3.4-fold higher in this SNP sparse region than in SNP dense regions C and D. Since we are estimating recombination frequency with a small number of crossovers, a bootstrap resampling method generates 95% confidence intervals based on 10,000 random iterations of our data, which shows SNP dense regions generally have lower recombination rates compared to the SNP sparse region than expected by random chance.

**Figure 5.**
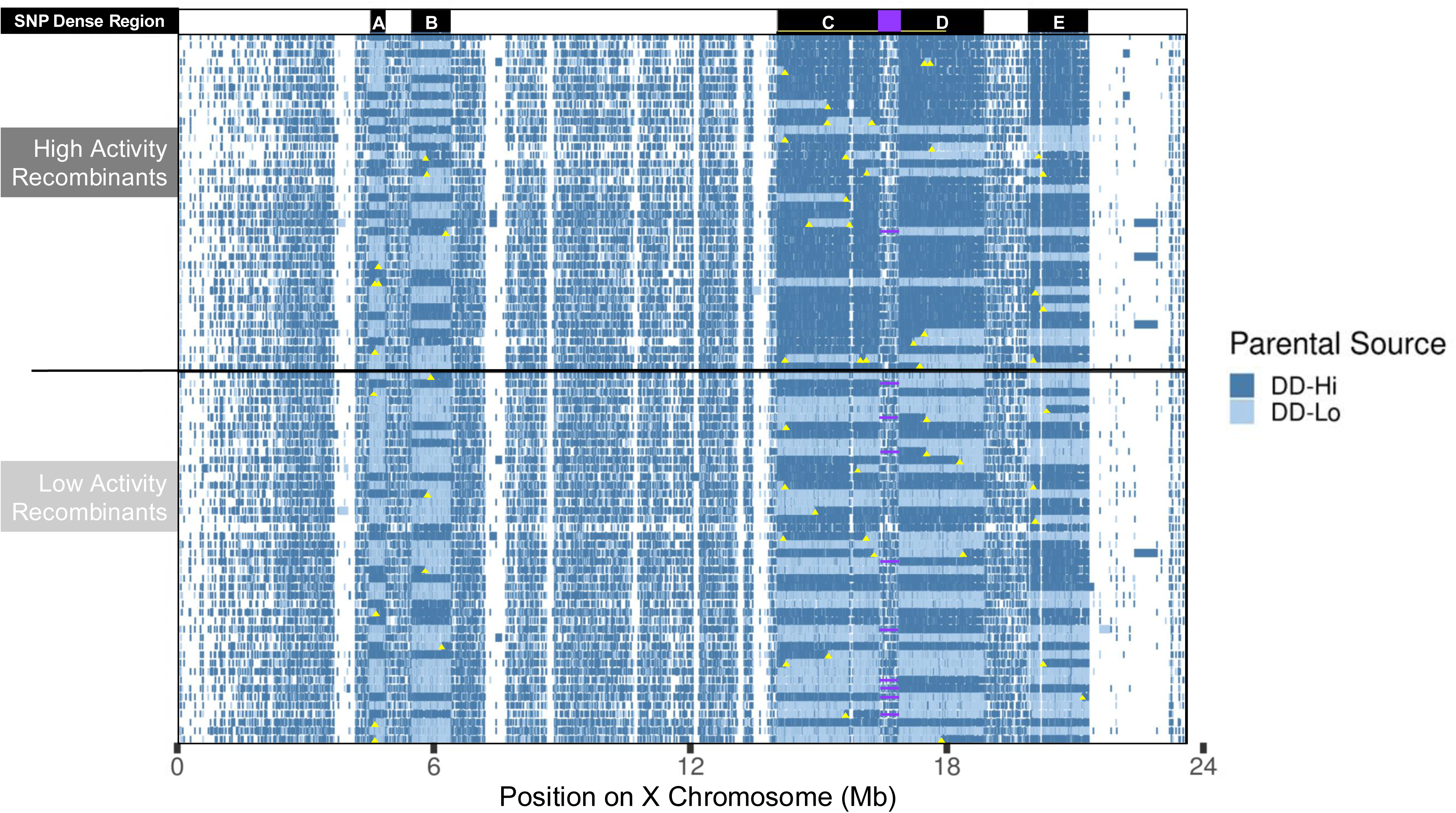
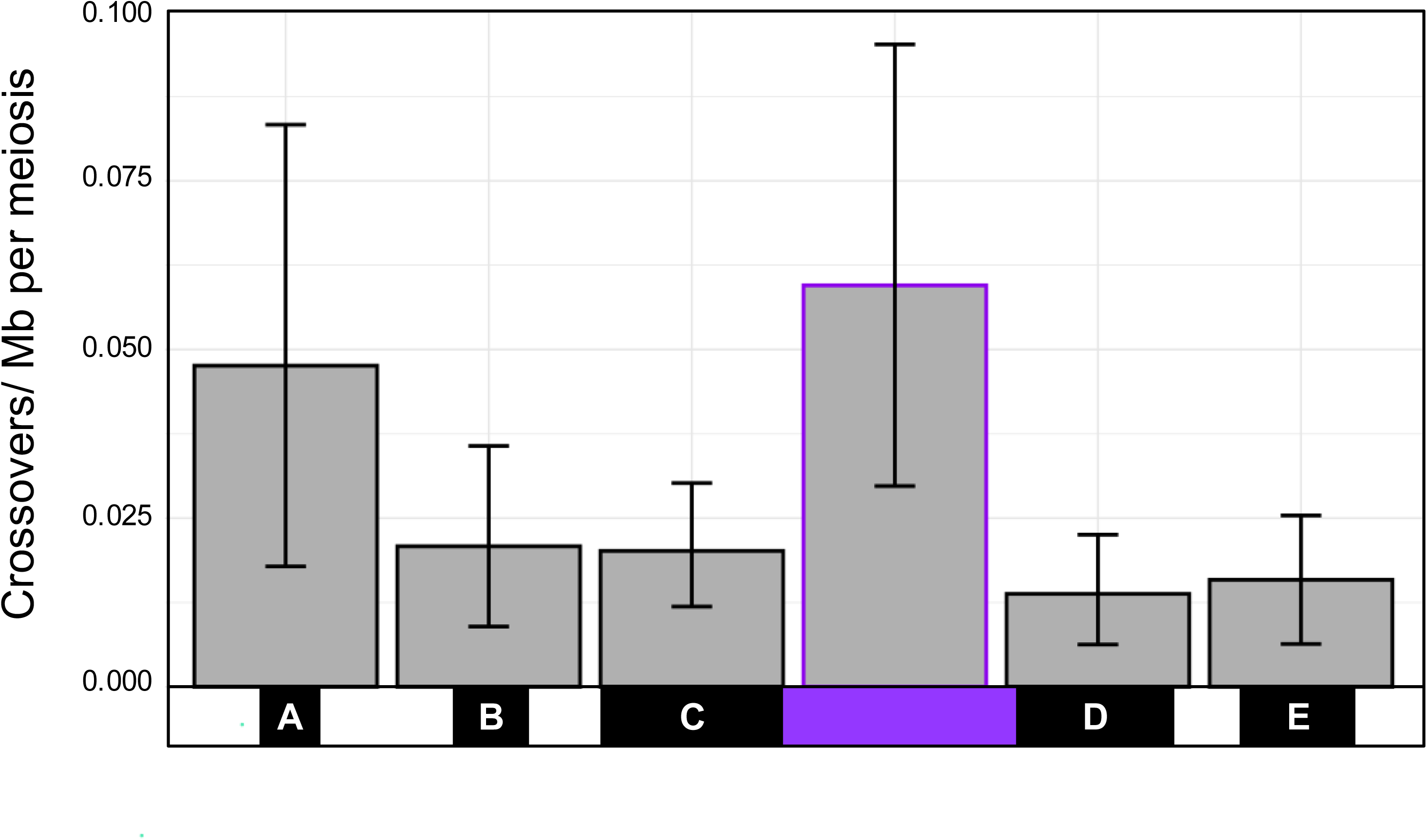
Recombination frequency within SNP dense regions is lower than in an adjoining SNP sparse region. The recombination frequency of each region was calculated for a single meiosis (see Methods). Crossovers per Mb was plotted with 95% confidence intervals generated using bootstrap resampling (n=10,00). SNP dense regions are denoted by black boxes that correspond to the length of each region; the smallest SNP sparse region used for comparison is denoted by the purple box.

### SNPs in regions C and D associate with activity levels in the recombinant flies

Our genetic mapping study identifies SNP dense regions C and part of D as containing single nucleotide differences associating with activity levels in the recombinant flies (Figure 3B). Casual observation of the data in Figure 4A appears to show much higher *DD-Hi* haplotype blocks in the high activity recombinants, and *DD-Lo* predominating in the low activity recombinants, especially in SNP dense regions C and D. We quantitated these parentage distributions in Figure 6, showing fractional *DD-Hi* parentage for each fly, plotting the mean fraction of parentage for the 40 high and 44 low activity recombinants in each SNP dense region. The distribution of individual recombinants is shown with a violin plot, since the majority of the data is either 0 (full *DD-Lo*) or 1 (full *DD-Hi)*. Parentage of the SNP dense regions that overlap the regions of high association show *DD-Hi* vs *DD-Lo* parentage that strongly correlates with activity levels of the recombinant flies, reaching significance in the 13.9-16.5 Mb region C. The correlation of parentage in the 16.9-18.8 Mb region D is similarly biased, but not nearly as significant as region C, presumably since only part of this region shows significant association. These observations are consistent with the importance of these associating regions in controlling the locomotor activity phenotype.

**Figure 6.**
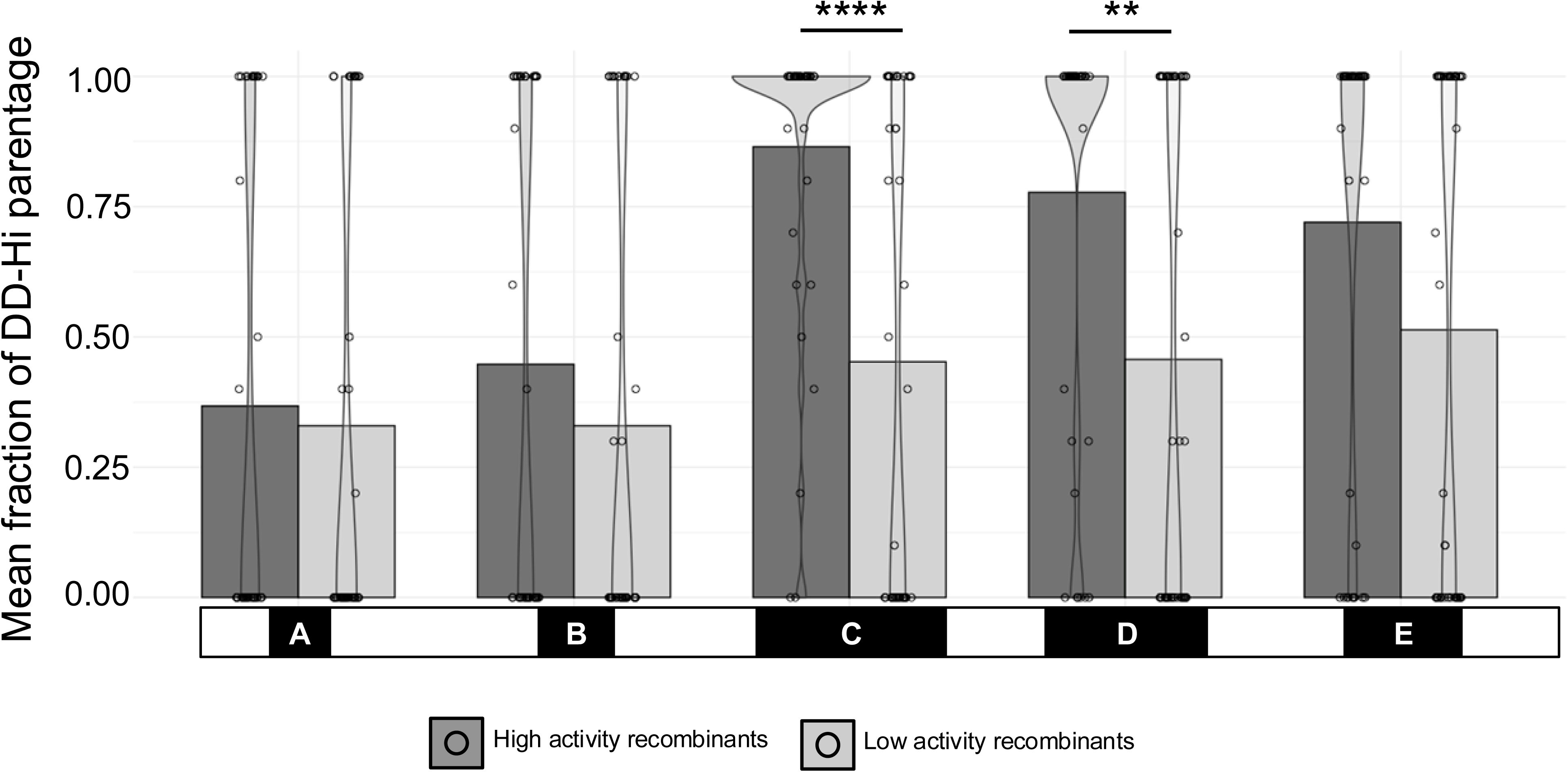
Locomotor activity of each recombinant line plotted as a function of the ratio of *DD-Hi* parentage within each SNP Dense region. The percentage of *DD-Hi* parentage was estimated within each SNP dense region for each recombinant fly by counting a full *DD-Hi* tract as 1 and a full *DD-Lo* tract as 0; recombinant flies that contained a crossover event were estimated as a fraction of *DD-Hi* parentage. Data presented as mean with the distribution of individual data points shown by the overlaying violin plot. Asterisks denote significant differences between the high and low activity recombinants within a given region where *\*\*p < 0.01 and ****p < 0.0001* [Wilcoxon Rank Sum Test - Bonferroni Corrected, n=40 high activity and 44 low activity].

### Uniform recombination mapping in earliest studies

As summarized above, a number of studies show variation in recombination rates across the central region of the *Drosophila* X-chromosome. Given the results we present here, we suspect that these studies could have been influenced by variable distributions of SNPs across the chromosomes. As such, we wondered whether the earliest mapping studies, all of which apparently used the same fly background, would be free from these mapping influences from distinct fly backgrounds. We took the X-chromosome mapping data from (Morgan et al., 1925), and plotted the given map position in Centimorgans versus the molecular position of the respective genes in flybase.org. These original mapping data were only corrected for frequencies of double recombinants. As shown in Figure 7, the genes map to a line with an r^2^ value of >0.99. Thus, this original map data is incredibly precise, strongly suggesting that later studies were inadvertently affected by inhomogeneous SNPs between the lines used in the crosses.

**Figure 7.**
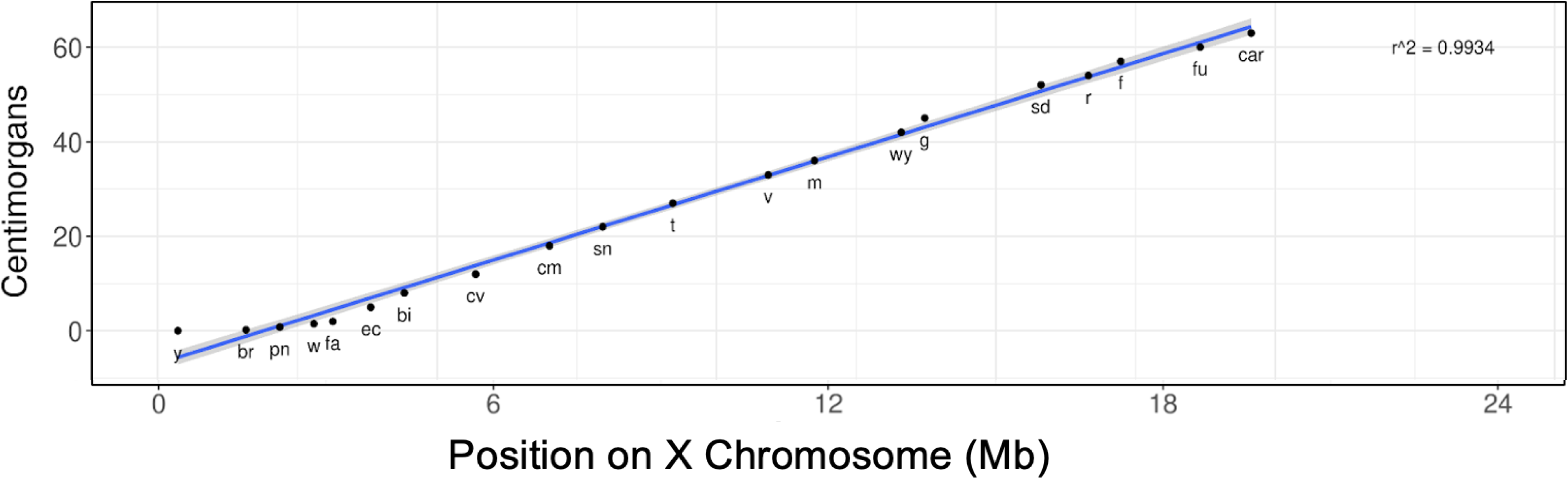
Recombination mapping of classical alleles. Recombination mapping is highly accurate across the central region of the X chromosome when assaying classical alleles on the X-chromosome. Each point represents a visible gene, mapped by both classical and molecular techniques. Recombination map positions and base positions on X-chromosome are from https://FlyBase.org. The datapoints map to a line with r2 =0.9974, plotting map position in centimorgans vs molecular position on the X-chromosome.

## Discussion

Dopamine is a critical regulator of arousal and locomotor activity in *Drosophila*. Flies lacking brain dopamine exhibit the expected phenotype of diminished locomotor activity (Cichewicz et al., 2017). Remarkably, in this study, we identify a sub-line of dopamine-deficient flies that recover normal levels of locomotor activity, despite the complete absence of brain dopamine. Using a non-bulked segregant approach and whole genome sequencing, we mapped the genetic basis of this recovery to a ∼3 Mb region of the X-chromosome. However, further fine mapping was hindered by an uneven distribution of single nucleotide differences within the starting X-chromosomes, biasing recombination events and thus limiting our resolution.

While our mapping focused on the X chromosome, we also detected minor associations with the high activity phenotype on the second chromosome. However, due to the nature of our sequencing approach, which used single males with diploid autosomes, we were unable to distinguish individual autosomes. This limitation made it difficult to confidently track inheritance patterns or pinpoint crossovers outside of the X chromosome. Given these constraints, we concentrated our analysis where we had the highest resolution.

The X-chromosomes used in our recombination study display an unusual pattern of sequence divergence, with regions of near-homozygosity, 0.04% divergence, interspersed with five distinct ∼1-3 Mbase regions with 0.25% average divergence. Our mapping identifies two of these SNP-dense regions, C and D (Figure 3), as associated with the high locomotor activity phenotype. However, the numbers of recombination events in these regions are lower than anticipated, at least partly accounting for the ∼5 fold lower resolution in our mapping versus the anticipated value. Counts of reciprocal crossovers confirmed a 3-fold reduction in crossover frequency relative to a nearby region of near homology, suggesting a novel mechanism limiting recombination in these SNP-dense regions. We note the possibility that non-random matings during the 15 generations during which free recombination occurred could have reduced the opportunity for effective recombination (Korol et al., 2000). However, our finding that the recombination frequency in the SNP-sparse region between the SNP-dense blocks C&D is at the expected frequency makes this unlikely.

Previous evidence in *Drosophila* shows variation in recombination rates across the *Drosophila* genome (Chan et al., 2012), particularly near the centromeric and telomeric regions where crossovers are strongly reduced (Comeron et al., 2012; Hughes et al., 2018; Lindsley & Sandler, 1977). Lindsley and Sandler (1977), show variation in the level of exchange across the X chromosome, but less extreme variation than on the autosomes. We note that the hot/cold spots seen by (Chan et al., 2012; Comeron et al., 2012) show variability between each study, and even between specific crosses within a given study (Comeron et al., 2012). We suggest that this variability could be related to patterns of varying blocks SNP density between starting strains as we have seen in this manuscript. The exception to this variation in recombination frequency across the X chromosome is seen in the earliest mapping studies from Morgan’s lab (Morgan et al., 1925). These early studies occurred at a time prior to the appearance of discrete background strains of *Drosophila,* which would presumably have differential patterns of SNPs.

We note that the SNP dense region C in our current data is less informative in our genetic mapping than we would have hoped. Figure 4A shows that high and low activity recombinants can have parentage mismatches from *DD-Hi* and *DD-Lo* on either side of these COs. These findings could be interpreted as showing that a large region is required to control activity levels. But this interpretation is confounded by the possibility of non-crossover gene conversion events that are beyond the detection level of our sequencing. This could explain our unexpected high activity recombinants that contain region C and D with total parentage from the *DD-Lo* parents, and vice versa.

Where is the genetic diversity in our starting lines coming from? The high and low activity dopamine deficient lines were isolated after we performed single pair matings of our dopamine deficient flies (Cichewicz et al., 2017) to our laboratory stock of *w^1118^*. We performed these crosses in an attempt to reduce genetic diversity in our dopamine deficient flies, but we suspect that instead we added genetic diversity. Our working stock of *w^1118^* has been maintained for decades, while at the same time we maintained a number of *w^1118^* stocks in diverse genetic backgrounds in our laboratory stock collection. Notes contemporaneous to when the crosses were performed indicate that our stock collection contained *w^1118^* in backgrounds including Canton-S, Oregon-R, Berlin, as well as *w^1118^* lines obtained from several colleagues around the world. When these lines were assayed for locomotor activity, they showed a 2-3 variation in locomotor activity (data not shown). We thus suspect that sometime in the past decades that our working stock of *w^1118^* was inadvertently contaminated with at least some of these genetic backgrounds. This hypothesis would help to explain the near identity along the majority X-chromosomes when comparing the *DD-Lo* and *DD-Hi* flies, since they would have been inter-mating prior to our single pair matings. The reduced crossover frequency in the punctuated regions of SNP density would thus serve to preserve these regions intact, consistent with our observations.

We find no direct precedent in *Drosophila* for our finding that reciprocal crossovers are selectively suppressed in chromosomal regions with enhanced divergence, though there are relevant examples in mitotic recombination and in other organisms. (Rutherford & Carpenter, 1988) examined isogenized X-chromosomes for differences in meiotic crossover frequencies relative to control chromosomes and found no differences, but the level of divergence in their control chromosomes was unknown. More recent studies show that pre-crossover enzymes bind less effectively to diverged regions during mitosis. In *Drosophila* cultured cells, bloom helicase binding and mitotic recombination is impeded by sequence divergence less than 0.5% (Kappeler et al., 2008), similar to the levels in our SNP dense regions. In mouse, divergence as high as 1.8% is not sufficient to trigger heteroduplex rejection during meiosis (Peterson et al., 2020). In yeast, it has long been appreciated that there is a reduced rate of meiotic recombination in heterozygous regions (Borts & Haber, 1987), and there is less binding of the mismatch repair protein Msh5 to regions of higher divergence (Dash et al., 2024).

Our findings underscore the importance of chromosomal relatedness when designing genetic mapping studies. Regions of punctuated diversity, like those identified in this study, can significantly skew recombination patterns, leading to biased mapping outcomes. This insight is critical for the broader field of genetic mapping in *Drosophila* and other organisms where similar recombination biases might occur unnoticed.

## Methods

### Measurement of locomotor activity

Locomotor activity was measured using the Drosophila Activity Monitor System (DAM, Trikinetics, MA USA) at 22C, 60% RH. Age matched flies were loaded into 3 mm ID glass tubes with standard cornmeal food on one end, and a cotton plug on the other. Entrainment was in a 12:12 light/dark cycle for 5 days, at which point they were switched to constant darkness, and activity was measured for an additional 5 days.

### Feeding of 3IY and Methamphetamine

3- Iodo-L-tyrosine (3-IY) (I8250, Sigma) and L-DOPA (D9628, Sigma) were mixed directly into melted fly food at final concentrations of 1.25 mg/ml and 1 mg/ml, respectively was dissolved in 1 ml of water at the 10X final concentration and mixed with 9 ml of melted fly food. The Trikinetics activity monitor tubes were loaded with a plug of food at one end, with drug feeding throughout the course of the experiment.

### HPLC analysis of dopamine and serotonin

Conditions were as per (Hardie & Hirsh, 2006), with a mobile phase consisting of 12% Acetonitrile, 0.4 mM Decanesulfonic acid, 50 mM sodium citrate/acetate, pH 4.5, 0.1 mM EDTA.

### Interbreeding *DD-Hi and DD-Lo* for Non-Bulked Segregant Analysis

*DD-Hi* and *DD-Lo* flies were allowed to freely intermate for 15 generations. The study started with group of ∼80 *DD-Hi* females and ∼55 *DD-Lo* males that were allowed to mate and lay eggs for five days in 3.07 L Tupperware containers with standard cornmeal media and yeast, at 22°C, 60% RH. Parental lines were cleared as soon as progeny eclosed. Progeny were transferred to fresh Tupperware containers to continue interbreeding with a transferred population of several hundred. After 15 generations, 788 males were collected for assay of locomotor activity as above. Forty-five of the least active and most active flies, representing activity levels of 95-238 and 1460-1861, respectively, were selected and DNA was isolated for sequencing. Additionally, three *DD-Hi* and *DD-Lo* females were sequenced for analysis of parentage.

### Sequencing

Libraries were constructed for paired-end Illumina sequencing in a HiSeq X. DNA isolation from whole flies was performed using an alcohol precipitation protocol adapted from (Bergland et al., 2014). Fragmentation, end repair, dA-tailing, and size selection was then achieved using the [NEBNext® Ultra II FS DNA Library Prep Kit for Illumina](https://www.neb.com/products/e7805-nebnext-ultra-ii-fs-dna-library-prep-kit-for-illumina%23Product%25252520Information), (NE Biolabs, Beverly, MA). Indexing primers for multiplexing were subsequently ligated using [NEBNext® Multiplex Oligos for Illumina® (Dual Index Primers Set 1)](https://www.neb.com/products/e7600-nebnext-multiplex-oligos-for-illumina-dual-index-primers-set-1%23Product%25252520Information) (NE Biolabs, Beverly, MA). We performed paired-end sequencing, generating approximately 4–6 million reads per sample with a read length of 150 bp, and removed adapter contamination using fastp. Of the 45 recombinant male flies sequenced, 40 high activity recombinant and 44 low activity recombinant sequences passed quality control and were used for downstream analysis. Of the three *DD-Hi* and three *DD-Lo* parental female flies sequenced, three *DD-Hi* and two *DD-Lo* sequences passed quality control and were used for downstream lineage analysis.

### SNP calling

SNPs were called using a combination of bioinformatic tools:

fastp was used in paired end mode and input 5’-AGATCGGAAGAGCACACGTCTGAACTCCAGTCA-3 as the adapter sequence for read 1 and 5’-GATCGGAAGAGCGTCGTGTAGGGAAAGAGTGT-3 ’as the adapter sequence for read 2 (Chen et al., 2018). Default fastp flags for quality control and trimming were used.

Alignment to the *Drosophila* genome assembly release 6 plus ISO1 MT (GCF_000001215.4) was performed using the BWA mem algorithm in paired end mode (Li & Durbin, 2009). Alignment flags included -M for downstream Picard compatibility and -a to allow for all output alignments. BAM files were sorted by read name using samtools for duplicates to be marked in the next step (Danecek et al., 2021). Duplicate reads were removed using MarkDuplicates from Broad Institute’s Picard toolkit (https://broadinstitute.github.io/picard/) flagging REMOVE_DUPLICATES as true. BAM files were then coordinate sorted using samtools, which is required for downstream analysis. Following alignment and duplicate removal, the Germline short variant discovery (SNPs + Indels) pipeline was followed using the Broad Institute’s Genome Analysis Toolkit (GATK) (https://gatk.broadinstitute.org/hc/en-us). The following GATK commands were used: HaplotypeCaller called variants on each sample’s BAM file and used the flag -ERC GVCF for reference model calling, which is necessary for multi-sample variant calling; GenomicsDBImport imported multiple gVCF files into a GenomicsDB workspace; and GenotypeGVCF called genotypes of gVCF files from a database that included parental data. GATK variant calling for 84 samples used – sample-ploidy 1 and 5 parental samples used –sample-ploidy 2.

In cases where SNPs are not detected at given base locations, it is inferred that these base pairs share the same origin as the nearest base pair for which a SNP was successfully called and matched to a parental genotype.

### Association Analysis

The variant call file generated from the GATK pipeline was analyzed for statistical association using PLINK v1.9.0-b.7.8 (Chang et al., 2015). For association analysis, we removed parental samples from combined vcf file and generated vcf file containing the only 84 male samples. We used the following arguments as inputs to PLINK along with the phenotype file and the 84 male samples vcf file as inputs.

--assoc fisher --allow-extra-chr --allow-no-sex --pheno phenotype_file1.txt

P values were generated using the Fisher’s exact test option. Manhattan plots were generated in R using ggplot2 (Wickham, 2016).

### Lineage Analysis

Parental identity was identified using the vcf generated from Broad Institute’s GATK pipeline. Each sample was evaluated at each SNP to determine if it matched uniquely to a parental line. SNPs for which a given sample had no supporting reads were ignored. SNPs matching any of the three parental samples were assigned a source lineage only if the genotype in the parental samples differed between *DD-Hi* and *DD-Lo*. Next, each sample was evaluated using the locations for which we had explicit sequencing data. Base pair locations which were missing sequencing data were interpolated to be the same as the surrounding SNPs. Ambiguous regions are indicated as white in Figure 4.

### Determining fraction of *DD-Hi* parentage

*DD-Hi* parentage was estimated within each SNP dense region by assigning either 0 for a full *DD-Lo* tract, 1 for a full *DD-Hi* tract, or a fraction representing the amount of *DD-Hi* for each fly. The mean values for high and low activity recombinants in each region were plotted using ggplot2 in R with the distribution of parentage values shown by the violin plots (Wickham, 2016). P-values were generated using a Wilcoxon Rank Sum test followed by Bonferroni correction for multiple comparisons.

### Estimating recombination frequency

Recombination frequency was estimated by counting changes in parental identity within the SNP dense regions, or between the smallest SNP sparse region (Figure 5). The total number of crossovers for a given region was divided by the length of the region to normalize to numbers of crossovers per 1 Mb. This was divided by 84 (total number of flies) and then 5 (total number of generations with recombination occurring) to quantify the recombination frequencies for a single meiosis within in each region. Bootstrap resampling (n = 10,000) was then used to generate 95% confidence intervals for each region.

### Long read sequencing to detect X-chromosome inversions

Nanopore long read sequencing was used to search for inversions, with sequence coverage of 23.4x for *DD-Lo* and 17.5x for *DD-Hi,* respectively. Structural variant analysis was performed using SVIM, which outputs all possible structural variants, even ones that have minimal read support. SWIM outputted no inversions with strong support. *DD-Lo* was found to have at most 29 inversions with very weak support with low numbers of supporting reads in the single digits, and at most 20 inversions with very weak support in *DD-Hi.* Only 0.41% and 0.43% of the X chromosome has 0 coverage for DD-Lo and DD-Hi, respectively, as measured by samtools depth. Therefore, we conclude there are no X-chromosome inversions inhibiting recombination in our fly strains.

## Acknowledgments

We acknowledge productive use of the fly genome database http://flybase.org. Clay Ford from the UVA StatLab provided critical code and advice for the bootstrap resampling analysis. We appreciate helpful comments from Alan Bergland, Ben Blackman, Eric Alani, Katja Kasimatas, Amanda Gibson, and helpful suggestions on the manuscript from Scott Hawley. We acknowledge support from the University of Virginia Bioinformatics Core, RRID:SCR_012718.

## Data Availability

Raw whole genome sequencing data is deposited in the NCBI SRA (http://www.ncbi.nlm.nih.gov/sra) as a BioProject under accession number PRJNA1393392.

## References

Andretic, R., van Swinderen, B., & Greenspan, R. J. (2005). Dopaminergic modulation of arousal in Drosophila. Current Biology: CB, 15(13), 1165–1175. 10.1016/j.cub.2005.05.025

Bergland, A. O., Behrman, E. L., O’Brien, K. R., Schmidt, P. S., & Petrov, D. A. (2014). Genomic evidence of rapid and stable adaptive oscillations over seasonal time scales in Drosophila. PLoS Genetics, 10(11), e1004775. 10.1371/journal.pgen.1004775

Birman, S., Morgan, B., Anzivino, M., & Hirsh, J. (1994). A novel and major isoform of tyrosine hydroxylase in Drosophila is generated by alternative RNA processing. The Journal of Biological Chemistry, 269(42), 26559–26567.

Borts, R. H., & Haber, J. E. (1987). Meiotic recombination in yeast: Alteration by multiple heterozygosities. Science (New York, N.Y.), 237(4821), 1459–1465. 10.1126/science.2820060

Chan, A. H., Jenkins, P. A., & Song, Y. S. (2012). Genome-wide fine-scale recombination rate variation in Drosophila melanogaster. PLoS Genetics, 8(12), e1003090. 10.1371/journal.pgen.1003090

Chang, C. C., Chow, C. C., Tellier, L. C., Vattikuti, S., Purcell, S. M., & Lee, J. J. (2015). Second-generation PLINK: Rising to the challenge of larger and richer datasets. GigaScience, 4, 7. 10.1186/s13742-015-0047-8

Chen, S., Zhou, Y., Chen, Y., & Gu, J. (2018). fastp: An ultra-fast all-in-one FASTQ preprocessor. Bioinformatics (Oxford, England), 34(17), i884–i890. 10.1093/bioinformatics/bty560

Cichewicz, K., Garren, E. J., Adiele, C., Aso, Y., Wang, Z., Wu, M., Birman, S., Rubin, G. M., & Hirsh, J. (2017). A new brain dopamine-deficient Drosophila and its pharmacological and genetic rescue. Genes, Brain, and Behavior, 16(3), 394–403. 10.1111/gbb.12353

Comeron, J. M., Ratnappan, R., & Bailin, S. (2012). The many landscapes of recombination in Drosophila melanogaster. PLoS Genetics, 8(10), e1002905. 10.1371/journal.pgen.1002905

Danecek, P., Bonfield, J. K., Liddle, J., Marshall, J., Ohan, V., Pollard, M. O., Whitwham, A., Keane, T., McCarthy, S. A., Davies, R. M., & Li, H. (2021). Twelve years of SAMtools and BCFtools. GigaScience, 10(2), giab008. 10.1093/gigascience/giab008

Dash, S., Joshi, S., Pankajam, A. V., Shinohara, A., & Nishant, K. T. (2024). Heterozygosity alters Msh5 binding to meiotic chromosomes in the baker’s yeast. Genetics, 226(3), iyad214. 10.1093/genetics/iyad214

Friggi-Grelin, F., Coulom, H., Meller, M., Gomez, D., Hirsh, J., & Birman, S. (2003). Targeted gene expression in Drosophila dopaminergic cells using regulatory sequences from tyrosine hydroxylase. Journal of Neurobiology, 54(4), 618–627. 10.1002/neu.10185

Hardie, S. L., & Hirsh, J. (2006). An improved method for the separation and detection of biogenic amines in adult Drosophila brain extracts by high performance liquid chromatography. Journal of Neuroscience Methods, 153(2), 243–249. 10.1016/j.jneumeth.2005.11.001

Hoffmann, A. A., & Rieseberg, L. H. (2008). Revisiting the Impact of Inversions in Evolution: From Population Genetic Markers to Drivers of Adaptive Shifts and Speciation? Annual Review of Ecology, Evolution, and Systematics, 39, 21–42. 10.1146/annurev.ecolsys.39.110707.173532

Hughes, S. E., Miller, D. E., Miller, A. L., & Hawley, R. S. (2018). Female Meiosis: Synapsis, Recombination, and Segregation in Drosophila melanogaster. Genetics, 208(3), 875–908. 10.1534/genetics.117.300081

Jones, G., Kleckner, N., & Zickler, D. (2024). Meiosis through three centuries. Chromosoma, 133(2), 93–115. 10.1007/s00412-024-00822-0

Kappeler, M., Kranz, E., Woolcock, K., Georgiev, O., & Schaffner, W. (2008). Drosophila bloom helicase maintains genome integrity by inhibiting recombination between divergent DNA sequences. Nucleic Acids Research, 36(21), 6907–6917. 10.1093/nar/gkn793

Korol, A., Rashkovetsky, E., Iliadi, K., Michalak, P., Ronin, Y., & Nevo, E. (2000). Nonrandom mating in Drosophila melanogaster laboratory populations derived from closely adjacent ecologically contrasting slopes at “Evolution Canyon.” Proceedings of the National Academy of Sciences of the United States of America, 97(23), 12637–12642. 10.1073/pnas.220041397

Li, H., & Durbin, R. (2009). Fast and accurate short read alignment with Burrows-Wheeler transform. Bioinformatics (Oxford, England), 25(14), 1754–1760. 10.1093/bioinformatics/btp324

Lima, S. Q., & Miesenböck, G. (2005). Remote control of behavior through genetically targeted photostimulation of neurons. Cell, 121(1), 141–152. 10.1016/j.cell.2005.02.004

Lindsley, D. L., & Sandler, L. (1977). The genetic analysis of meiosis in female Drosophila melanogaster. Philosophical Transactions of the Royal Society of London. Series B, Biological Sciences, 277(955), 295–312. 10.1098/rstb.1977.0019

Mao, Z., Bozzella, M., Seluanov, A., & Gorbunova, V. (2008). Comparison of nonhomologous end joining and homologous recombination in human cells. DNA Repair, 7(10), 1765–1771. 10.1016/j.dnarep.2008.06.018

Michelmore, R. W., Paran, I., & Kesseli, R. V. (1991). Identification of markers linked to disease-resistance genes by bulked segregant analysis: A rapid method to detect markers in specific genomic regions by using segregating populations. Proceedings of the National Academy of Sciences of the United States of America, 88(21), 9828–9832. 10.1073/pnas.88.21.9828

Miller, D. E., Takeo, S., Nandanan, K., Paulson, A., Gogol, M. M., Noll, A. C., Perera, A. G., Walton, K. N., Gilliland, W. D., Li, H., Staehling, K. K., Blumenstiel, J. P., & Hawley, R. S. (2012). A Whole-Chromosome Analysis of Meiotic Recombination in Drosophila melanogaster. G3 (Bethesda, Md.), 2(2), 249–260. 10.1534/g3.111.001396

Morgan, T. H., Bridges, C. B., & Sturtevant, A. H. (1925). Linkage. In The Genetics of Drosophila (pp. 87–108).

Nestler, E. J. (2004). Historical review: Molecular and cellular mechanisms of opiate and cocaine addiction. Trends in Pharmacological Sciences, 25(4), 210–218. 10.1016/j.tips.2004.02.005

Peterson, S. E., Keeney, S., & Jasin, M. (2020). Mechanistic Insight into Crossing over during Mouse Meiosis. Molecular Cell, 78(6), 1252–1263.e3. 10.1016/j.molcel.2020.04.009

Riemensperger, T., Isabel, G., Coulom, H., Neuser, K., Seugnet, L., Kume, K., Iché-Torres, M., Cassar, M., Strauss, R., Preat, T., Hirsh, J., & Birman, S. (2011). Behavioral consequences of dopamine deficiency in the Drosophila central nervous system. Proceedings of the National Academy of Sciences of the United States of America, 108(2), 834–839. 10.1073/pnas.1010930108

Rutherford, S. L., & Carpenter, A. T. (1988). The effect of sequence homozygosity on the frequency of X-chromosomal exchange in Drosophila melanogaster females. Genetics, 120(3), 725–732. 10.1093/genetics/120.3.725

Wickham, H. (with Sievert, C.). (2016). *ggplot2: Elegant graphics for data analysis* (Second edition). Springer international publishing.

Yellman, C., Tao, H., He, B., & Hirsh, J. (1997). Conserved and sexually dimorphic behavioral responses to biogenic amines in decapitated Drosophila. Proceedings of the National Academy of Sciences of the United States of America, 94(8), 4131–4136. 10.1073/pnas.94.8.4131

